# A Novel Algorithm for the Harmonization of Pan-cancer Proteomics

**DOI:** 10.1101/2025.03.17.642820

**Authors:** Nitzan Simchi, Ofer Givton, Joseph Rinberg, Alon Shtrikman, Tamar Geiger, Alexey I. Nesvizhskii, Eran Seger, Kirill Pevzner

**Affiliations:** Protai Bio, Ramat Gan, Israel; Weizmann Institute of Science, Rehovot, Israel; Department of Computational Medicine and Bioinformatics, University of Michigan, Ann Arbor, MI 48109, USA

## Abstract

Proteomic characterization of cancer tissues holds the potential to advance therapeutic options and reveal novel biomarkers by unlocking insights available only on the proteome level. However, proteomics data analysis is greatly challenged by systematic technical variability in experimental protocols, instrumentation and data processing, restricting comparisons between studies. With the continued and unprecedented growth of proteomics datasets, a comprehensive strategy for harmonizing these datasets is necessary to enable large-scale integrative analyses. Herein, we describe a novel framework for pan-cancer harmonization and imputation, which offers the scientific community an updated approach to this challenge. Rather than relying on a single batch-effect correction algorithm, our multi-step approach accurately addresses critical systematic differences with custom-tailored solutions, including standardized reanalysis of raw data and an autoencoder for pan-cancer integration. By introducing a suite of benchmarks, we bridge the critical gap in reliable harmonization evaluation. Using this framework, we created a harmonized pan-cancer dataset and demonstrated its superiority over existing solutions and previous pan-cancer harmonization efforts. We further demonstrated its utility in revealing prognostic markers, estimating indication-wide biomarker prevalence, and facilitating target discovery for cancer subtypes. We expect our work to provide powerful tools supporting proteomics research for precision cancer medicine.

## Introduction

High-throughput proteomics has revolutionized cancer research by enabling the comprehensive profiling of protein expression across diverse tumor types. Tandem Mass Tag (TMT) labeling, in particular, allows for the simultaneous quantification of proteins in multiple samples, facilitating large-scale comparative studies. However, integrating proteomic data from different studies poses significant challenges due to batch effects—systematic technical variations arising from differences in experimental protocols, instrumentation, and data processing pipelines. These batch effects can obscure true biological signals, hinder cross-study comparisons, and impede the discovery of novel biomarkers and therapeutic targets.

Batch effects in proteomics stem from various sources, including discrepancies in sample procurement or preparation methods, variations in mass spectrometry settings, and the use of different data analysis algorithms^1^. For instance, inconsistencies in protein extraction procedures or peptide identification criteria can lead to significant differences in protein quantification between studies.^2^ Additionally, the use of varied protein databases and normalization techniques exacerbates these discrepancies, making it difficult to discern whether observed differences in protein expression are due to biological variability or technical biases. This issue is particularly pronounced when integrating datasets generated across different laboratories, each employing distinct protocols and analytical approaches.^3^

Traditional statistical methods for batch effect correction, such as ComBat^4^ and Limma^5^, have been applied to mitigate technical variability in proteomic data. While these methods can adjust for certain types of batch effects, they often assume linear, additive relationships and require complete data matrices without missing values. Proteomic datasets are inherently sparse due to the stochastic nature of peptide detection in mass spectrometry, and often exhibit batch-specific patterns of missing values that complicate batch correction. Such batch-dependent missingness complicates downstream imputation, which may itself be influenced by the underlying batch effects. Moreover, these methods may not adequately capture complex, nonlinear batch effects or preserve intricate biological relationships within the data, such as protein–protein interactions and pathway activities.

The challenge of batch effect correction is further compounded in pan-cancer analyses encompassing multiple tissue types where often a dataset would analyze single indication and tissue type, creating complete confoundment between tissue and dataset-of-origin. Biological variability between different cancer indications introduces additional complexity, as batch correction methods must distinguish between true biological differences and technical artifacts. Preserving meaningful biological signals while eliminating batch effects is crucial for accurate downstream analyses, including the identification of cancer subtypes and exploration of disease mechanisms. Failure to adequately address batch effects can lead to misleading conclusions, such as false associations or the overlooking of clinically significant biomarkers.

Previous efforts to integrate proteomic datasets across studies have highlighted both the potential benefits and limitations of existing approaches. The Clinical Proteomic Tumor Analysis Consortium (CPTAC)^6^ has generated extensive proteomic data for various cancer types, providing valuable resources for the research community. However, residual batch effects and inconsistencies in data processing methodologies still impede comprehensive analyses and cross-study comparisons.^3^ These limitations underscore the pressing need for robust harmonization strategies capable of effectively integrating heterogeneous proteomic datasets while preserving the integrity of biological information.

Addressing these challenges requires a multifaceted approach that considers both technical and biological sources of variability. Standardizing data processing pipelines is a critical first step. Reanalyzing raw data using consistent parameters and protein databases can minimize variability introduced during data processing and enhance comparability across studies. This includes uniform protocols for peak detection, peptide identification, and protein quantification. By controlling these variables, researchers can reduce technical discrepancies and focus on genuine biological differences.

Advanced computational methods also play a pivotal role in effective data harmonization. Machine learning techniques, such as autoencoders^7^ and other neural network architectures, offer promising avenues for modeling complex, nonlinear relationships inherent in proteomic data. These methods can learn to disentangle technical artifacts from true biological signals by capturing underlying patterns in high-dimensional datasets. Additionally, innovative approaches to handle missing data are essential, as traditional imputation methods may introduce biases or fail to account for the intricacies of proteomic measurements.

Here, we developed a novel harmonization framework designed to integrate proteomic data from multiple studies into a cohesive pan-cancer protein expression atlas. By leveraging standardized data reanalysis, advanced normalization techniques, and machine learning approaches, we effectively mitigated batch effects while preserving meaningful biological variability. This integrated dataset can facilitate more accurate cross-study comparisons, enhance statistical power for detecting subtle biological signals, and support downstream analyses such as biomarker discovery and therapeutic target identification.

A key distinguishing feature of our approach is the application of autoencoders (AEs) to amplify biological signals while effectively reducing noise in proteomics datasets. In addition, we introduce a novel imputation algorithm that accurately preserves tissue-specific signals, while simultaneously eliminating batch effects that occur when comparing datasets from the same tissue. Unlike existing methods, our algorithm continuously improves with the addition of more samples and datasets per tissue, demonstrating its scalability and adaptability. Using a novel, biologically-driven harmonization evaluation pipeline, our results show a marked improvement over previous approaches, both in terms of accuracy and consistency, highlighting the algorithm’s ability to enhance data harmonization in a field where comparability between studies is often a major hurdle. Combining these together we supply an end-to-end framework for harmonization and benchmarking, thereby improving data comparability across studies.

## Results

### Algorithm overview and development process

To support the development of our harmonization algorithm, we initially collected and curated a diverse set of publicly available datasets, including both Formalin Fixed Paraffin Embedded (FFPE) tissue samples and Fresh-Frozen (FF) bulk tumor tissue samples. Data included tumors from multiple types of cancer indications, including breast cancer (BRCA), epithelial ovarian cancer (EOC), uterine corpus endometrial carcinoma (UCEC), pancreatic ductal adenocarcinoma (PDAC), colorectal cancer (CRC), hepatocellular carcinoma (HCC), glioblastoma (GBM), lung adenocarcinoma (LUAD), lung squamous cell carcinoma (LSCC), clear cell renal cell carcinoma (CCRCC), and head & neck squamous cell Carcinoma (HNSCC). We decided to focus on a single type of data generation technique, namely Tandem Mass Tag (TMT) labeling, due to the relative popularity of high quality TMT datasets.

Our harmonization algorithm follows a multi-step approach, each tackling one aspect, and designed to effectively integrate data from multiple proteomics datasets, addressing the challenges posed by varying experimental protocols, instrumentation, and data processing pipelines (Fig. 1). Below, we provide a step-by-step description of the algorithm and the key techniques utilized at each stage.

**Figure 1.**
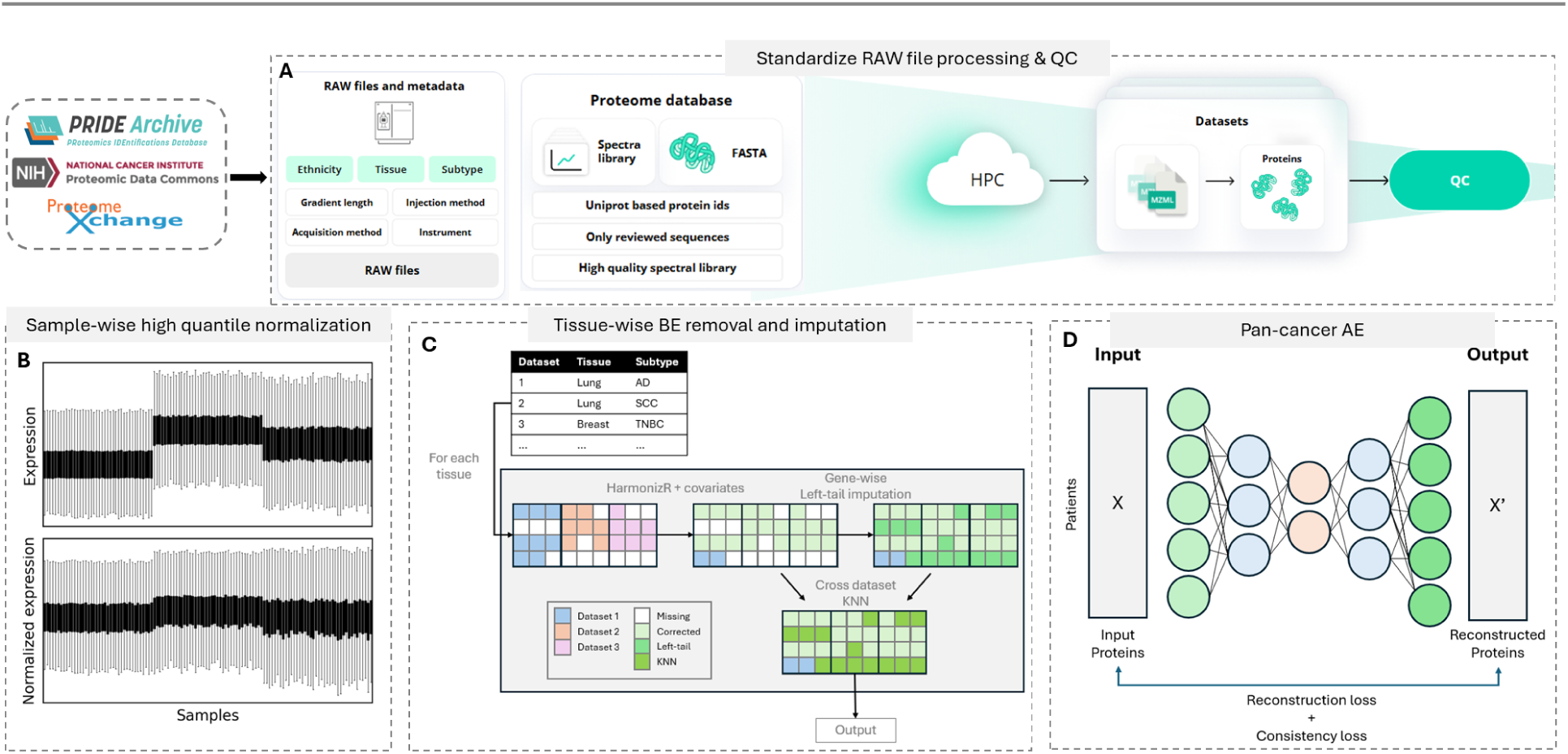
Pan-cancer proteomics harmonization algorithm. Our multi-step approach tackles different aspects of batch effects in proteomics data. Beginning with standardized reanalysis of raw data (A), we then proceed to sample-wise normalization (B), followed by indication-wise imputation coupled with batch-effect removal (C), and concludes with a pan-cancer autoencoder for biological signal smoothing (D).

**Figure 2.**
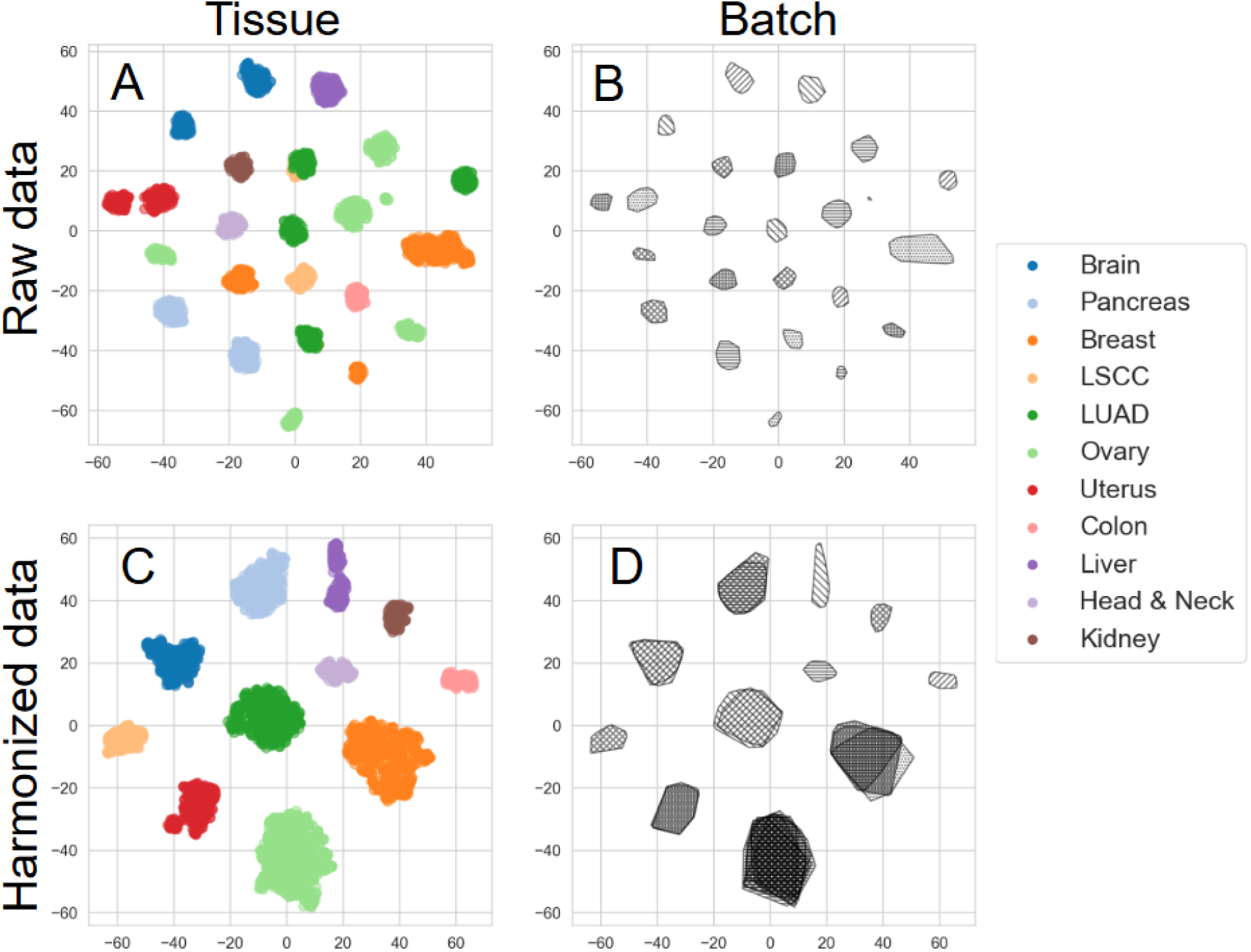
Batch effect improvements after running harmonization. Performing t-SNE on the expression data and coloring/hatching by tissue/batch highlights the batch effects visually. Although each tissue type clusters together in the raw data (A), marking the same plot by batch shows that this is simply because each run was of a single tissue type (B). After harmonization, tissues cluster together strongly (C) and are no longer confounded by the batch (D).

#### Step 1: Standardized Reanalysis of RAW Data

Unlike batch effects arising from experimental conditions or sample preparation variability, the data processing pipeline, a major source of batch effects, can be controlled. Therefore, the first crucial step in our algorithm is the standardized reanalysis of RAW data. The necessity for reanalysis arises from the significant variability found in published proteomics datasets^35^. Many studies use inconsistent protein databases such as UniProt or RefSeq, often including isoforms or unreviewed sequences. Worse still, more often than not, different studies utilize different data normalization techniques (e.g., z-score, or ratio data for TMT labeling), making protein quantification completely dataset-relative. These inconsistencies have a dramatic effect on harmonization efforts, effectively rendering published, processed TMT datasets “unharmonizable”. By reanalyzing the RAW data, we can ensure that the final integrated dataset is coherent and comparable, facilitating downstream analyses. Additionally, using a smaller, more consistent search space reduces the false discovery rate (FDR) and improves the reliability of results across studies.

First, we identified the optimal parameters for integrating batches from various studies, including peak detection, peptide identification, and protein quantification, which are key to maximizing the consistency of results across different platforms and methods. In parallel with the reanalysis, we implement a stringent quality control (QC) process to ensure the integrity of the integrated dataset. Briefly, we excluded low-confidence peptides to reduce noise in the final dataset, while requiring a minimum number of peptides per protein to ensure consistent quantification. Additionally, we required minimal coverage of proteins on a sample-wise basis. Datasets where this requirement wasn’t met by any sample were excluded entirely. Full details describing the parameters and thresholds used in the reanalysis process are provided in the Methods.

Processing such vast amounts of data demanded scalable computational resources. We employed a custom-made cloud-based infrastructure, similar to QuantMS^8^, enabling dynamic expansion through continuous reanalysis and integration of new datasets as they become available, without compromising computational efficiency. Taking advantage of recent advances in proteomic data processing algorithmic pipelines (e.g. FragPipe^9^), compared to the published processed data, we were able to improve data quality and coverage without altering the FDR threshold (See Methods and Supplementary Table).

#### Step 2: Sample-wise high quantile normalization

Sample-wise normalization can manage, to some extent, the significant variability of the measurement scale and protein quantity between samples, particularly across datasets^10^. Unlike median centering, a common practice which normalizes based on the central tendency, high quantile (HQ) normalization focuses on normalizing the higher end of the intensity distribution. Intensities in the higher quantiles are typically measured with greater precision, making them a reliable anchor for normalization. We tested HQ scaling against several sample-wise and protein-wise scaling through our harmonization evaluation pipeline, with HQ scaling outperforming all other scaling methods (Table 4).

#### Step 3: Tissue-wise batch effect removal and imputation

Missing values pose a significant challenge for harmonization. Not only do different batches carry distinct missing value patterns, effectively making them a fundamental batch effect by themselves, most batch effect correction methods cannot handle missing values, requiring imputation prior to correction. To solve this problem, by leveraging information from other datasets of the same tissue types to impute missing values, we developed a novel algorithm, seeking to turn this setback into an opportunity for better harmonization.

First, we applied HarmonizR^11^, a recently developed tool that employs matrix dissection to apply ComBat across datasets without requiring prior imputation of missing values. As ComBat assumes that datasets originate from the same underlying distribution, and because different cancer tissues pose the first major source of biological variability in our pan-cancer atlas, we applied our imputation algorithm in a tissue-wise manner. However, even within a single cancer type this assumption is often violated by significant heterogeneity arising from distinct subpopulations. To accommodate this, we extended HarmonizR to incorporate covariates, such as PAM50 subtypes in breast cancer, allowing us to better preserve subtype-specific signals (Supplementary Fig. 2).

Next, we performed protein-wise left-tail imputation, drawing from a lower quantile of the distributions, to generate an imputed reference matrix. Using this complete reference matrix, we performed T-SNE dimensionality reduction to enhance local, multidimensional similarities, and found the K-nearest neighbors for each sample in the reduced space. Finally, for each sample, we imputed missing values by the K-nearest neighbors in *other* datasets only, replacing the left-tail estimates. By imputing data between batches, we ensure unbiased imputation, disentangling the batch effects from the missing values pattern.

Importantly, looking at the samples in a lower dimensional space, this step generated perfect clustering of samples by their respective tissues, without significant clustering by dataset of origin.

#### Step 4: Pan-cancer AutoEncoder

So far, we have attempted to minimize batch effects globally through standardized reanalysis and scaling, and specifically mitigated batch effects in a tissue wise manner through ComBat. However, when dealing with complex batch effects confounded with biological variables, ComBat’s assumptions of linear, additive and univariate batch effects, may result in improper corrections or the removal of true biological signals. Furthermore, using ComBat in a tissue-wise manner may bias cross-tissue differences, limiting pan-cancer analyses.

To tackle these limitations, we employed an autoencoder model, a neural network architecture excelling at compressing high-dimensional data into a lower-dimensional representation, or “latent space,” and then reconstructing the original data from this compressed form. By compressing data from different cancer types into a unified latent space, the model identifies common biological signals that transcend individual datasets and tissue types. This helps amplify real biological signals while reducing dataset-specific noise and batch effects that may remain even after initial correction.

The training process was guided by minimizing two losses; a reconstruction loss, which ensures that the model accurately reconstructs the input data from its compressed representation. Additionally, we introduce a “*consistency loss”* that penalizes the model for deviations from the original protein-wise expression ranking of samples within each dataset separately, as we assume that within each dataset, the relative sample expression is generally correct. Working in tandem, reconstruction loss ensures that the autoencoder captures the key underlying biological features necessary to reconstruct the input, while, on the other hand, consistency loss encourages the model to maintain the correct relative expression patterns of proteins within a dataset.

On the tissue level, the autoencoder dramatically improved the biological signal, coupled with a significant improvement in the similarity of identical samples (Table 3). We speculate that with the introduction of more datasets for the autoencoder training in the future will allow for a more successful and accurate harmonization.

### Biologically-driven benchmarks

Biologically driven benchmarks are crucial for evaluating the effectiveness of harmonization methods. Assessing the preservation of known biological signals, such as tissue-specific markers or protein complexes, provides insight into whether harmonization efforts are maintaining data integrity. Housekeeping genes, expected to exhibit stable expression across samples, serve as negative controls to ensure that batch correction methods are not introducing artificial variability. Furthermore, analyzing cross-batch replicates—identical samples processed in different batches—can directly measure the success of batch effect removal.

Data harmonization efforts may cause either incomplete removal of batch effects, or the removal of true biological signals. Owing to their complex nature, evaluating the residual batch effects presents a major challenge, intertwined with the harmonization process itself and crucial to its success. Therefore, harmonization evaluation played a pivotal role in our work. We assumed successfully harmonized data should bring samples from different batches to a shared multidimensional space, while preserving the underlying biological relationships between samples and proteins within each individual batch. To that end, we applied a series of harmonization benchmarks, aimed at capturing different biological ground truths of the integrated dataset, thus guiding the development of our algorithm. Because different harmonization strategies offer trade-offs in performance, our goal was to build an algorithm that balances all tradeoffs.

To evaluate the biological variability between cancer tissues, we analyzed the differential expression of tissue-markers. Tissue-markers are proteins assumed to be over-expressed in specific tissues or cancer indications, thus providing a positive control of inter-tissue differences. It is of high importance to note that the termed tissue-markers were not aimed to be *tissue-specific* markers, i.e. proteins expressed only in specific tissues, but universally abundant proteins expressed across many tissues. That is because this benchmark is meant to capture more delicate inter-tissue differences, whereas tissue-specific markers would most likely cause missing values for these markers in other tissues, therefore leading any cross-tissue analysis of such markers to be highly dependent on their imputed values. The tissue-markers curation process used two pan-cancer proteomics datasets^12,13^, where tumor samples from different tissues were analyzed together. Then, we selected proteins that met our selection criteria in both datasets. Finally, we filtered the selected tissue-markers for those validated in literature (see Methods and Supplementary Table).

For the benchmark, we first calculated the fold change for each tissue marker between samples of its target tissue and all other tissues. Then, we repeated this calculation for 1,000 randomly selected protein-tissue pairs, and took the 90th percentile of the random distribution as a threshold for differential tissue expression. Finally, we determined the percentage of the tissue-markers with a fold-change above our threshold. Additionally, we performed a Mann-Whitney U-test between the two distributions with the alternative hypothesis that the tissue-markers tend to be higher than the random markers for their respective tissues (Fig. 3A).

**Figure 3.**
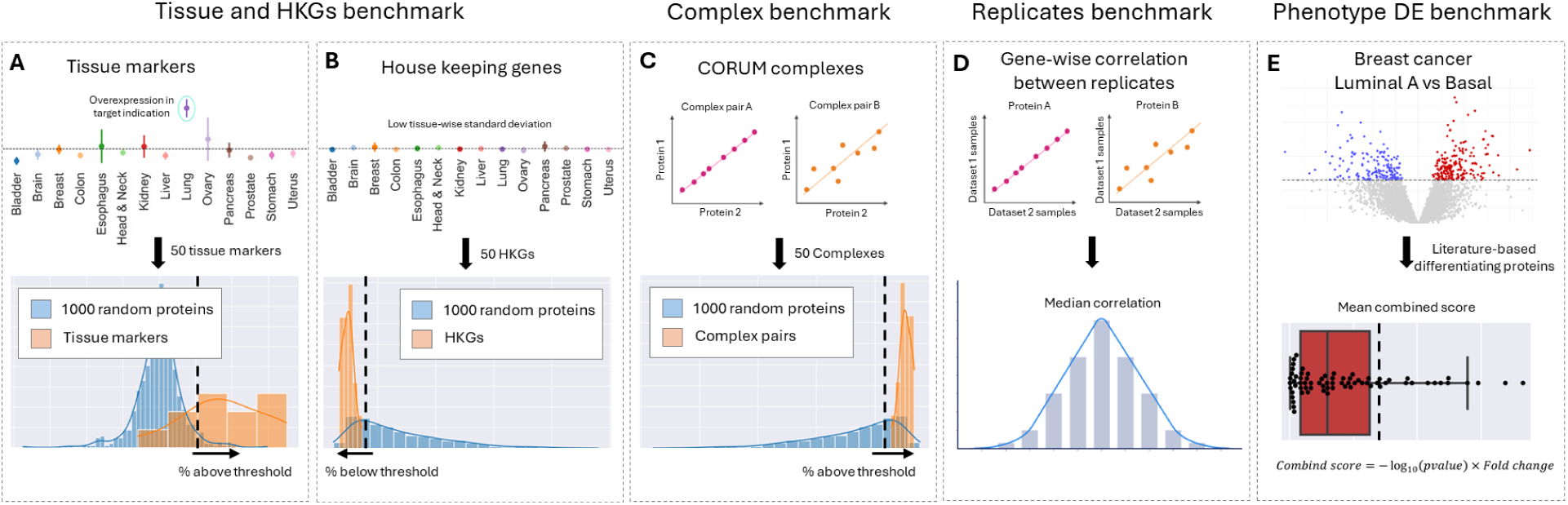
Biologically-driven benchmarks description. The benchmark results are illustrated by the following plots: (A) For tissue markers, we display one protein example that is overexpressed in the desired indication and the distribution of the 50 markers compared to 1000 random markers. (B) Similarly, the housekeeping genes benchmark is shown. (C) For the complex benchmark, we present a histogram of the complex pairs correlation compared to random pairs. (D) For the replicates benchmark, a histogram of the gene-wise correlation between replicates is shown where we take the median correlation. (E) Lastly, for the phenotype differential expression benchmark, we show the combined score of the known differentiating proteins.

The housekeeping genes benchmark is conceptually similar to the tissue-markers, but serves as a negative control for tissue-specific signals. Opposite to tissue-markers, HKGs (housekeeping genes) are expected to exhibit minimal variability between tissues, providing a “mirror image” to the tissue-markers. The selection process and criteria for the HKGs were similar to that of the tissue-markers, with the only exception being the selection of proteins that were found to be least variable between tissues.

First, we calculated the tissue-weighted standard deviation for the HKGs, followed by a similar calculation for all other proteins in our data. We considered the 10th standard-deviation percentile of all proteins to be the threshold for minimally variable proteins, and determined the percentage of HKGs with a standard deviation below the threshold. Mann-Whitney U-test between the two distributions with the alternative hypothesis shows that the HKGs tend to be lower than all other proteins (Fig. 3B).

When considering protein expression data on an absolute scale, biological signals appear in two axes; first, protein-wise differences in protein expression between samples; and second, sample-wise differences between proteins. Many batch effect correction methods (e.g. ComBat^4^, HarmonizR^11^), use statistical modeling techniques to arrange protein-wise differences between samples of different batches on the same scale. However, such data manipulation risks damaging the sample-wise relationships between proteins. Additionally, due to inherent differences in mass-spectrometry peptide ionization potential, we cannot assume a linear relationship between the measured MS intensity and the absolute protein abundances within a sample. This nonlinear relationship can be significantly influenced by batch effects. Therefore, successful batch effect removal is expected to preserve or enhance the natural dependencies between proteins.

To evaluate the expected relationships between proteins within samples, we curated a list of PCPs (protein complex pairs), assumed to be exclusively co-expressed in specific protein complexes, thus exhibiting high correlation and similar expression values. We calculated the correlation coefficient and the absolute deviation between PCPs. We repeated the absolute deviation calculation for 1,000 randomly selected protein pairs, and performed a Mann-Whitney U-test between the two distributions and calculated the p-value with the alternative hypothesis that the absolute deviation between PCPs tends to be lower than that of random pairs. Additionally, we considered the percentage of PCPs with a correlation coefficient above 0.7 (Fig. 3C).

As a direct measurement of protein-level harmonization, we took advantage of a unique study design in two pairs of batches in our harmonized atlas (Hu et al.^14^ and Mcdermott et al.^15^; Chowdhury et al.^16^ Fresh-frozen and Formalin Fixed Paraffin Embedded batches). In these dataset pairs, technical replicates of the same tumor samples were analyzed in separate, distinct batches. To avoid noisy or flat proteins, we only considered the top 50% proteins with the highest interquartile range (IQR) within each batch. Then, we calculated the protein-wise correlation coefficient between the two sets of replicates, and took the mean of the two correlation medians as the final score (Fig. 3D).

To evaluate the phenotypic variability within cancer indications, we compared two well defined breast cancer subtypes, Luminal A and Basal-like. We performed a differential expression analysis for proteins known to be differentially expressed in either subtype, and calculated a combined score for each protein. (Fig 3E; See Methods).

We tested our algorithm against two other common batch effect correction algorithms, ComBat and Limma, performed in an indication-wise manner following a similar rationale (see Methods). Benchmark results are summarized in Table 1. Our algorithm outperformed the others in enhancing both the intra- and inter-tissue biological signals. Specifically, it saw 83% of tissue markers passing the threshold compared to the second best result of 67% (p-value X and Y, respectively). In the phenotypic DE, it achieved a median adjusted p-value of 6.5e-25 compared to 4.3e-05 with Indication-wise Limma. Notably, our algorithm successfully increased the mean cross-batch replicates correlation to 0.52, with the other two algorithms failing to show any improvement from the baseline. Importantly, our algorithm managed to preserve the data integrity, as reflected in the HKG and PCP benchmarks (75% and 94% of proteins passing the threshold, respectively).

**Table 1.** Benchmark of harmonization algorithms.

|  | Baseline | Indication-wise<br>ComBat | Indication-wise<br>Limma | Autoencoder |
| --- | --- | --- | --- | --- |
| Tissue markers<br>(% above threshold) | 58% | 67% | 67% | 83% |
| Housekeeping genes<br>(% below threshold) | 69% | 63% | 75% | 75% |
| Protein complex pairs<br>(% above threshold) | 100% | 100% | 97% | 94% |
| Luminal A vs Basal DE<br>(mean combined score) | 2.0 | 13.1 | 11.6 | 25.3 |
| Luminal A vs Basal DE<br>(median adj. p-value) | 1 | 1e-03 | 4.3e-05 | 6.5e-25 |
| Replicates<br>(Mean correlation) | 0.29 | 0.3 | 0.29 | 0.52 |
| KNN-Batch model<br>(Classification accuracy) | 1.00 | 0.74 | 0.71 | 0.72 |

To date and to the best of our knowledge, few efforts to harmonize pan-cancer proteomics data have been made. We benchmarked the Clinical Proteomic Tumor Analysis Consortium (CPTAC) pan-cancer datasets and compared the results to our algorithm to two other harmonization studies by Li et al.^2^ and Zhang et al.^1^, in either case considering only the overlapping samples and proteins between our data and theirs (Table 2).

**Table 2.** Benchmark of harmonized datasets.

|  | Li et al.<sup>17</sup> | Autoencoder | Zhang et al.<sup>18</sup> | Autoencoder |
| --- | --- | --- | --- | --- |
| Tissue markers<br>(% above threshold) | 85% | 71% | 58% | 91% |
| Housekeeping genes<br>(% below threshold) | 21% | 78% | 18% | 81% |
| Protein complex pairs<br>(% above threshold) | 59% | 94% | 85% | 97% |

While the Li dataset was able to capture expected tissue-specific differences, slightly outperforming our harmonized atlas in the tissue-markers benchmark, this result came with a great cost, as only 21% of the HKGs passed the significance threshold. This result raises the suspicion that the Li dataset, aside from true tissue-specific variability, presents high false positive signals, pointing at an overall artificial pan-cancer variability. Looking at the PCP benchmark, where only 59% of PCPs passed the significance threshold, further shows how this dataset failed to preserve the underlying multidimensional protein relationships. Compared to the Zhang dataset, our harmonization achieved significantly better performance in all benchmarks. Notably, the Zhang dataset achieved poorer results in both the tissue-markers and HKG benchmarks, indicating that it failed to model the expected tissue-specific variability.

Finally, to demonstrate the justification for each step of the algorithm and its contribution, we present two final benchmarks. Table 3 displays the incremental performance gains achieved with each new step of the algorithm, while Table 4 offers a comparison of the various scaling methods.

**Table 3.** Contribution of harmonization steps.

|  | Reanalysis | Sample-wise HQ scaling | HQ scaling + Tissue-wise BE removal and imputation | Autoencoder |
| --- | --- | --- | --- | --- |
| Tissue markers (% above threshold) | 66% | 91% | 83% | 83% |
| Housekeeping genes (% below threshold) | 68% | 68% | 75% | 75% |
| Protein complex pairs (% above threshold) | 97% | 79% | 94% | 94% |
| Luminal A vs Basal DE (mean combined score) | 6.9 | 8.1 | 13.4 | 25.3 |
| Luminal A vs Basal DE (median adj. p-value) | 0.96 | 0.0049 | 9.6e-06 | 6.5e-25 |
| Replicates (Mean correlation) | 0.42 | 0.38 | 0.41 | 0.52 |

**Table 4.** Benchmark of scaling methods.

|  | Sample-wise high quantile scaling | Sample-wise robust scaling | Protein-wise robust scaling | Sample-wise median scaling | Protein-wise median scaling |
| --- | --- | --- | --- | --- | --- |
| Tissue markers (% above threshold) | 91% | 91% | 41% | 91% | 66% |
| Housekeeping genes (% below threshold) | 68% | 50% | 25% | 62% | 68% |
| Protein complex pairs (% above threshold) | 79% | 70% | 97% | 79% | 97% |
| Luminal A vs Basal DE (mean combined score) | 8.1 | 2.1 | 3.3 | 8.4 | 6.9 |
| Luminal A vs Basal DE (median adj. p-value) | 0.0049 | 0.0125 | 0.96 | 0.0071 | 0.96 |
| Replicates (Mean correlation) | 0.38 | 0.31 | 0.42 | 0.38 | 0.42 |

### Cross-Dataset Integration and Use Cases

#### Pan-Cancer Indication Expansion and Prioritization

A harmonized pan-cancer protein expression atlas serves as a powerful resource, overcoming limitations of smaller, indication-specific datasets. To demonstrate its potential in expanding and prioritizing therapeutic targets for cancer indications, we compared the mean tissue-weighted expression of therapeutic targets in their clinically approved indications (Fig. 4). When defining tissues with mean expression above the tissue-weighted median as clinically significant, our analysis achieved a recall of 0.72, demonstrating our harmonized atlas’ utility in accurately identifying clinically relevant indications. On the other hand, the relatively low precision of 0.35, when taken together with the relatively high recall, likely underscores the potential for clinical indication expansion, rather than misidentified target-tissue relationships.

**Figure 4.**
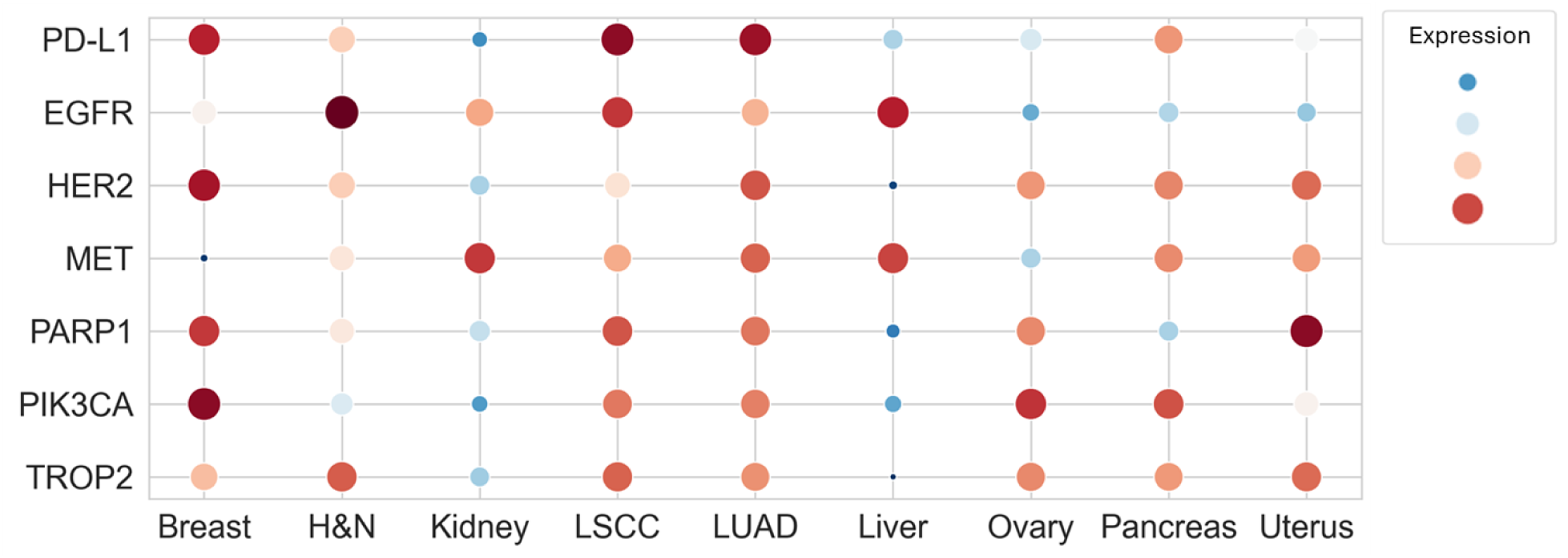
Pan-cancer biomarker expression prevalence. Z-scored expression of specific proteins across different indications. The larger the circle, the more the protein is expressed relative to the other proteins in the figure.

For example, the low expression of EGFR in pancreatic cancer compared to other cancers^19^, likely explains the limited success of EGFR inhibitors in clinical trials for this indication. While EGFR-targeted therapies combined with standard treatments like gemcitabine have been tested, they have shown minimal improvement in patient outcomes. This limited efficacy appears linked to the relatively low levels of EGFR expression in pancreatic tumors as seen in Fig. 4, reducing the therapeutic impact of EGFR inhibition and suggesting that alternative or combination treatment strategies may be required to achieve better clinical results.

Another interesting insight from Fig. 4 is the overexpression of MET and EGFR in hepatocellular carcinoma (HCC), which holds substantial potential for expanding therapeutic options, as these markers are increasingly implicated in promoting tumor progression and resistance to conventional treatments. EGFR inhibition, particularly when combined with other therapies like lenvatinib, an FDA-approved multikinase inhibitor for HCC, shows promising results in preclinical and early clinical studies. Lenvatinib alone offers limited efficacy, but when combined with EGFR inhibitors, it induces strong anti-tumor responses by targeting EGFR-PAK2-ERK5 signaling feedback loops, creating a synthetic lethality that limits tumor growth^20^. Moreover, EGFR inhibition may prevent HCC development by reducing liver fibrosis and metastasis, as well as overcome resistance in lenvatinib-refractory tumors. This therapeutic synergy not only underscores the significance of EGFR and MET as targets but also suggests their combined targeting could enhance treatment outcomes and prevent disease progression in liver cancer, warranting further investigation in larger clinical trials. In conclusion, pan-cancer protein expression can serve as a good predictor for clinical success.

#### Target Discovery for Pan-Cancer Subtypes

Although tumor protein expression is largely dominated by tissue-specific differences, various tumor types from different tissues share common activation of cancer-related pathways and aberrant signaling patterns. Therefore, by leveraging the harmonized pan-cancer dataset, we explored the existence of such pan-cancer subtypes. We performed Gene Set Variation Analysis (GSVA) across all samples, filtering for cancer-related pathways that were enriched in at least two tissue types. Hierarchical clustering of these data identified three main clusters (Fig. 5A); Cluster A, enriched in DNA damage repair and cell cycle regulation, was predominantly observed in ovarian, breast, uterine, LSCC and head & neck cancers. Cluster B, enriched in signaling and regulatory pathways in cancer progression, was mainly prevalent in LUAD and pancreatic cancers. Cluster C, enriched in metabolic pathways and energy production, prevalent in kidney and liver cancers. By repeating the analysis in the TCGA pan-cancer transcriptomics atlas, we were able to partially reproduce these findings, further supporting their validity (Fig. 5B). Differential mutation analysis revealed a significant enrichment of mutations relevant to specific clusters (Fig. 5C and Fig. 5E), such as mutations in EGFR and KRAS in Cluster B, or mutated PIK3CA in Cluster A (Chi^2^ test, p-values 1.5e-4, 4.6e-123 and 7.1e-15, respectively).

**Figure 5.**
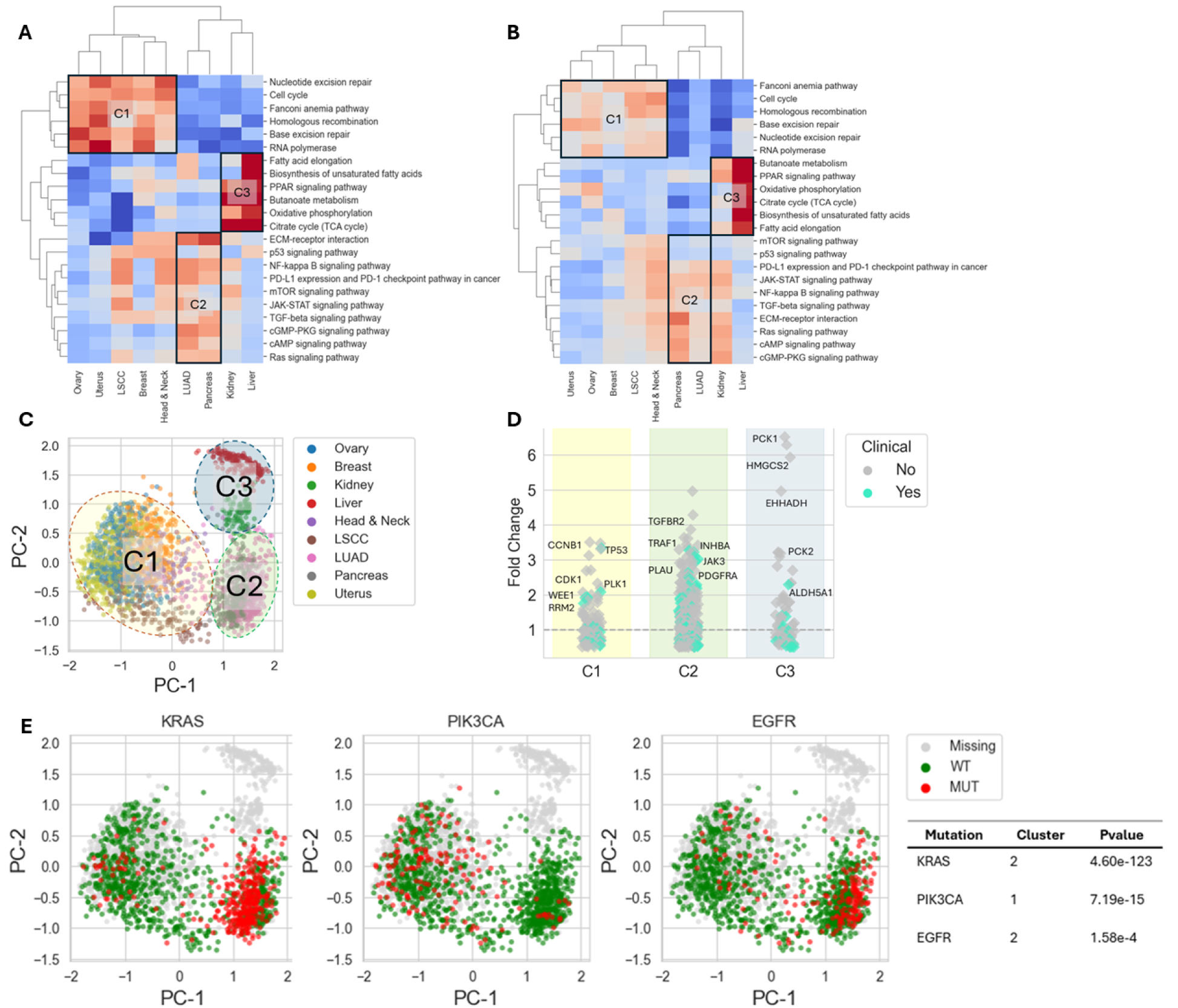
Pan-cancer therapeutic subtypes. (A) Hierarchical clustering of selected pathways enrichment scores using the harmonized pan-cancer data identifies 3 main clusters. (B) Similar analysis using TCGA pan-cancer transcriptomics atlas. (C) Sample level PCA using the harmonized pan-cancer data highlights the 3 clusters. (D) Differential expressed druggable targets within each cluster. (E) Differential mutation analysis highlights significant enrichment of mutations relevant to specific clusters.

Next, by looking at differentially expressed proteins within each cluster compared to the other ones, we identified druggable targets adhering to the clusters’ hypothesized mechanism driving tumorigenesis, with many already approved or in clinical trials (Fig. 5D). Well-known targets included CCNB1, CDK1, PLK1, and WEE1 in Cluster A, and JAK3, PDGFRA, VEGFA and PD-L1 in Cluster B. Additionally, we uncovered several novel potential therapeutic targets. BUB1B, SKP2, HMMR and UBE2T, involved in DNA repair and cell cycle regulation, were enriched in Cluster A. INHBA, a subunit of the activin protein involved in cell proliferation, differentiation and immune modulation signaling, was enriched in our “aberrant signaling” Cluster B. Cluster B further revealed potential new immunotherapy targets, such as ITGA2, ITGA9, CD247 and CXCL12, which are not currently addressed by any clinically approved drugs. HMGCS2 and PCK1, enzymes involved in ketogenesis and gluconeogenesis, respectively, were both enriched in our “metabolic” Cluster C.

Together, these findings underscore the ability of our harmonization approach to reveal subtypes that transcend traditional tissue boundaries, providing insights into shared oncogenic mechanisms across cancers, thus facilitating a more comprehensive understanding of cancer biology and enabling the identification of druggable targets that could be pursued across multiple cancers.

#### Consistent Prognostic Biomarker Discovery for Lung Adenocarcinoma

Finally, we explored how integrating multiple harmonized datasets of a single cancer indication can increase statistical power and enhance the ability to answer indication-specific questions, such as prognostic biomarker discovery.

We first combined two independent LUAD datasets (Lehtiö et al.^21^ and Gillette et al.^22^) as our biomarker discovery data (training set). We split the patient samples into two prognosis groups: OS > 3 years (better prognosis) and OS < 2 years (worse prognosis). As a measure of variability, we calculated the median absolute deviation for all proteins, and selected the top 5,000 proteins. To establish differentially expressed proteins, we performed a multiple ANOVA between the two groups, filtered for proteins with an absolute fold change > 0.5, and considered the 20 proteins with the lowest p-value for further analysis. To avoid selecting only highly positively or negatively correlated proteins for our biomarker, we split the remaining proteins into two groups, positively and negatively correlated to the most differentially expressed protein, SLC2A1. Splitting our data into training and validation sets, we trained a logistic regression classifier on the train set, in each trial a combination of 1-4 proteins from each of the two differentially expressed protein groups (Fig. 6A). Taking the model with the highest receiver operating characteristic (ROC) area under the curve (AUC; 0.7) on the validation set, we identified a 4-protein prognostic biomarker (Fig. 6C). Projecting our 4 feature biomarker on a PPI network reveals it is part of two modules (Fig. 6B). The first, represented by LCN2, COL12A1, and THBS1, is involved in tumor cell migration, invasion, angiogenesis and interactions with the immune system. The second module, represented by ADH1B, is positively correlated with prognosis. With roles in ethanol and retinol metabolism, decreased levels of ADH1B confer metabolic regulation within tumor cells, increasing reliance on glycolysis (Warburg effect), thus facilitating a favorable outcome. Therefore, this module represents down-regulated proteins whose normal function interferes with cancer progression.

**Figure 6.**
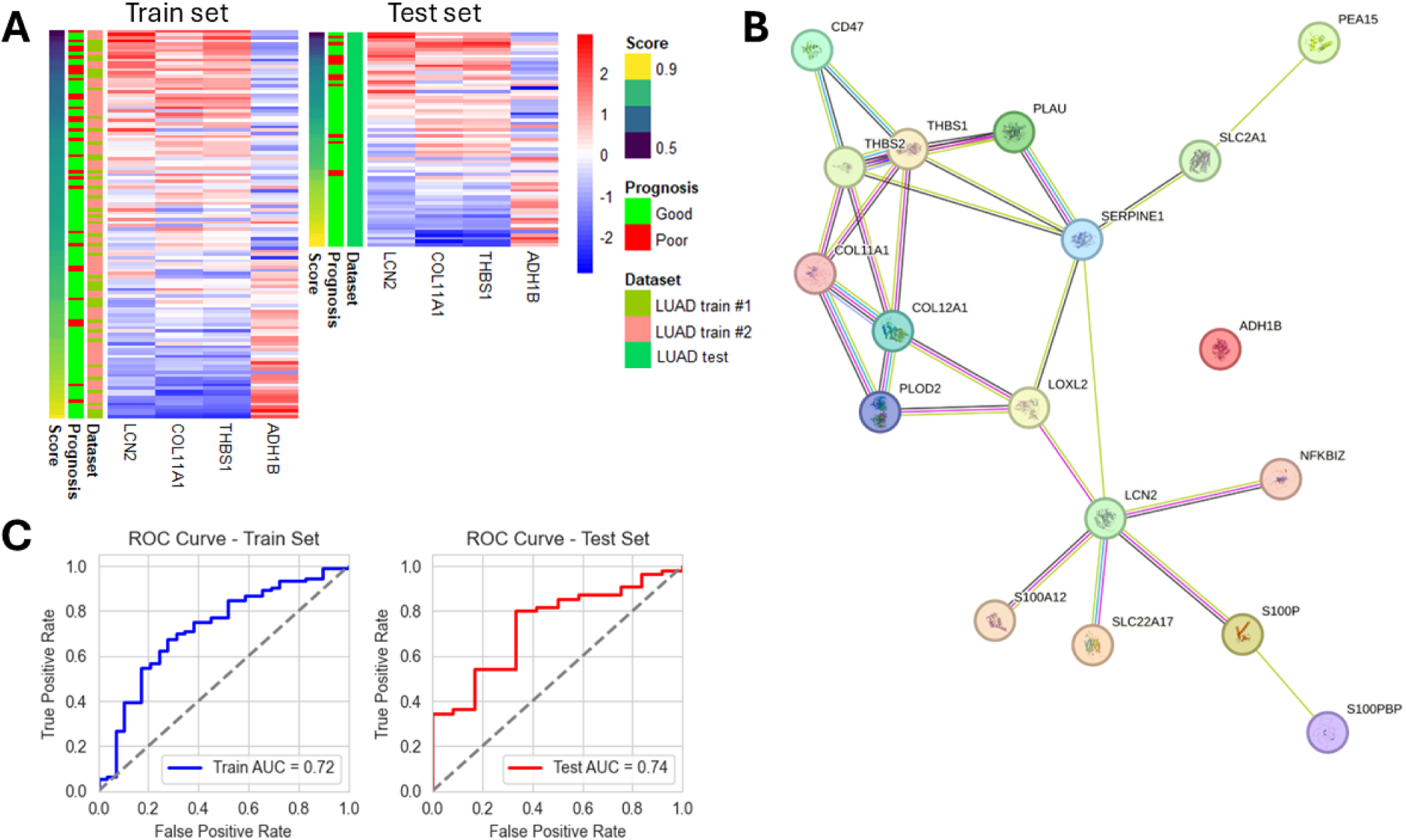
Lung Adenocarcinoma prognostic biomarker. (A) Hierarchical clustering of biomarker proteins for train and test sets. (B) PPI network of biomarker proteins. (C) Model performance on train and test sets.

To validate our findings, we applied our biomarker on a third, independent test LUAD dataset (Soltis et al.^23^), achieving an AUC of 0.76, superior to that of our training set (Fig. 6C). This process ensured that any identified prognostic markers were robust and generalizable across different LUAD cohorts. In conclusion, harmonization of different LUAD datasets with our algorithm enabled the discovery of a prognostic biomarker with mechanistic relevance of LUAD invasiveness, potentially offering novel therapeutic intervention points.

## Discussion

Batch effects pose a long-standing challenge in multi-study proteomics data integration, more-so in the setting of pan-cancer harmonization, where the majority of published datasets are confounded by sample tissue of origin. In this study, we present a novel, multi-step harmonization algorithm designed to address the unique characteristics of pan-cancer proteomics data, specifically TMT. Central to our approach is the idea that this is not a “one size fits all” problem, and no single batch effect correction method can cover all technical and biological sources of variability. Instead, we tackled these issues in stages, each tailored to address critical sources of variability; from standardized reanalysis of raw data to targeted, tissue-specific imputation, and finally an autoencoder-based correction designed to unify the entire proteomic landscape across cancer types.

One of the most critical insights we gained is the importance of starting with a clean foundation. By reanalyzing raw data using consistent databases and parameters - rather than relying on pre-processed, published datasets - we not only ensure coherent protein identification and quantification but also avoid inconsistencies that may be introduced when studies use different pipelines or normalization methods. When considering TMT data specifically, where proteome expression data is commonly published in the form of internally-normalized ratios to an internal standard, the reanalysis is a mandatory requirement for cross-study comparison. A potential downside of this approach is that reanalyzing a large number of publicly available datasets requires both significant computational resources and curation effort, which not all labs can afford. Interestingly, our sample-wise high-quantile scaling approach outperformed prevalent normalization methods. This observation should be validated in future studies.

Another pillar of our work was the realization that current tools and practices designed to evaluate batch-effect removal can be misleading, and rely heavily on strict presumption - whose violation is often overlooked. Therefore, we have developed a set of biological harmonization benchmarks, each specifically designed to provide positive and negative controls for each critical aspect or possible pitfall of harmonization. By utilizing these biological ground-truths, we can reliably test the strength of expected biological signals in the data and can, therefore, extrapolate that other, unknown natural phenomena and associations are also enhanced or preserved after the harmonization process. Preserving biological relationships while reducing artificial noise was central to the algorithm’s success. The tissue-marker and housekeeping gene benchmarks illustrated its capacity to maintain expected variability and stability in protein expression across cancer types. High correlation of protein complex pairs confirmed the algorithm’s ability to safeguard sample-wise protein relationships, a critical aspect of proteomics data integrity. Importantly, these achievements were realized without sacrificing data quality or introducing artifacts, as evidenced by consistent results across technical replicates and pan-cancer phenotypic analyses.

The inclusion of a pan-cancer autoencoder is vital. By compressing high-dimensional data into a unified latent space, the model identified common biological signals across different datasets and tissue types. This approach addressed limitations inherent in traditional linear methods, enhancing cross-batch consistency and amplifying true biological signals. The autoencoder’s ability to preserve inter-tissue biological differences while maintaining pan-cancer analytical capability resulted in significant improvements in intra-tissue harmonization compared to tissue-wise batch effect removal alone. It should be noted that this approach is limited by the amount of data available for training, including multiple datasets of each tissue type, which may not always be available.

Comparisons with existing harmonized proteomics datasets further emphasized the strengths of our approach. While the dataset by Li et al.^17^ performed well in capturing tissue-specific variability, it introduced significant false-positive signals, as reflected in poor housekeeping gene stability and protein complex pair correlation. In contrast, our harmonized dataset balanced biological signal retention with data integrity, achieving superior results in both tissue-specific and global benchmarks. This demonstrates the algorithm’s capacity to produce reliable, biologically relevant harmonized proteomics datasets. We highlight the translational potential of using a harmonized pan-cancer proteomics atlas for biomarker discovery.

Further directions to investigate consists of support for additional protocols such as LFQ and DIA, for which we suspect the Autoencoder will give a great benefit, as well as harmonization of post-translational modification data. Furthermore, a comprehensive analysis of the latent space representations generated by the Autoencoder model could potentially uncover novel insights that are specific to certain tissues or shared across multiple tissue types. This deeper investigation may reveal previously unknown relationships between protein expression patterns and tissue-specific functions, leading to a better understanding of the underlying biological mechanisms. Additionally, exploring the learned embeddings could identify common regulatory networks or signaling pathways that are conserved across different tissues, providing valuable information about fundamental biological processes. Lastly, the algorithm can be applied to cell lines datasets, as well as proteomics datasets outside cancer, depending on the amounts of data available.

In conclusion, a large scale pan-cancer harmonization algorithm is a valuable resource for the scientific community, enabling new discoveries in cancer biology, biomarker research, and therapeutic development. By addressing key challenges in harmonization, this work provides a powerful framework for advancing precision oncology research.

## Methods

### Data Description

To support the development of our pan-cancer harmonization pipeline, we curated multiple cancer proteomics datasets to establish a comprehensive resource of protein expression patterns across various cancer types. We prioritized contemporary datasets from recent years, with a large number of individual patient samples, high protein coverage depth, and as many datasets per individual cancer tissue. Generally, proteomic profiling can be achieved by label free-methods, with either data-dependent or data-independent acquisition (DDA and DIA, respectively), or with chemical labeling techniques, such as iTRAQ or Tandem Mass Tag (TMT). Because each of these methods presents an individual data modality with unique characteristics and biases, we chose to focus on one. As we found TMT datasets matching our selection criteria to be relatively more abundant, the TMT method was chosen for this study.

Raw data files were sourced from several publicly available repositories, including Proteomic Data Commons^24^, ProteomeXchange^25^, and PRIDE^26^. The data covered studies conducted between 2019 and 2024. The collected datasets encompassed proteomic profiles from 11 distinct cancer types: Breast cancer (BRCA), epithelial ovarian cancer (EOC), uterine corpus endometrial carcinoma (UCEC), pancreatic ductal adenocarcinoma (PDAC), colorectal cancer (CRC), hepatocellular carcinoma (HCC), glioblastoma (GBM), lung adenocarcinoma (LUAD), lung squamous cell carcinoma (LSCC), clear cell renal cell carcinoma (CCRCC), and head & neck squamous cell Carcinoma (HNSCC). The atlas included a total of 2789 FFPE and Fresh-Frozen tumor samples. Where available, matching RNA, mutations, and clinical annotations data were manually downloaded from respective papers’ Supplementary Materials.

To support the comparison of our harmonized atlas to other publicly available, pan-cancer harmonized datasets, we collected data from the studies of Li et al.^17^ and Zhang et al.^18^ Processed proteomic profiling data was downloaded from the papers’ Supplementary Materials.

### Tissue-markers curation process and benchmark

To evaluate the expected biological differences between cancer tissues, we analyzed the differential expression of tissue-markers. We defined tissue-markers as universally abundant proteins expressed across many tissues and differentially expressed in specific cancer tissues. We took advantage of the works by Zhou et al.^12^ and Song et al.^13^, who composed two “internally harmonized” pan-cancer datasets - i.e. single datasets where tumor samples from different tissues were analyzed together. First, we filtered the top 1000 most universally abundant proteins (proteins with least missing values) in our harmonization-naive pan-cancer atlas. Then, in each pan-cancer dataset, we further filtered for the top 10% highly expressed proteins. Then, in each dataset and for each indication, we performed a multiple one-sided t-test between samples belonging to the target indication and samples from all other indications and took the 10% proteins with the highest t-statistic as the most differentially expressed proteins for. Finally, for the resulting proteins in each indication, we evaluated the union between the two datasets, and further refined them through manual visual validation and a literature review of their association with the target tissue. In equivocal cases, we attempted to validate the results with the CCLE pan-cancer cell-line data or the Human Protein Atlas. The final resulting tissue markers are provided in Supplementary.

To score the tissue-markers benchmark, we calculated the fold change for each tissue marker, by subtracting its mean expression in the associated tissue from its mean expression in all other tissues. Then, we repeated this calculation for 10,000 randomly selected protein-tissue pairs and took the 90th percentile of the random distribution as a threshold for differential tissue expression. The percentage of tissue-markers passing this threshold served as this benchmark’s score. Additionally, to determine statistical significance, we performed a one-tailed Mann-Whitney U-test between the two resulting distributions with the alternative hypothesis that the tissue-markers tend to be higher than the random markers.

### Housekeeping Genes (HKGs) curation process and benchmark

To serve as a negative control for tissue-specific signals, we utilized Housekeeping Genes (HKGs), which are expected to show minimal variability between tissues. The curation process of the HKGs was conceptually similar to that of the tissue-markers, except instead of taking the most differentially expressed proteins between indications, calculated the mean tissue-wise expression of each protein, and considered the proteins with the lowest standard deviation of tissue-weighted means. The full list of curated HKGs is provided in Supplementary.

To score the HKG benchmark, we first calculated the tissue-weighted standard deviation for the HKGs, i.e. the standard deviation of the tissue-wise mean expression of each HKG. Then, we performed a similar calculation for all other proteins in our data. We considered the 10th standard-deviation percentile of the all-proteins distribution to be the threshold for minimally variable proteins, and determined the percentage of HKGs with a standard deviation below the threshold. To determine statistical significance, we performed a Mann-Whitney U-test between the two distributions with the alternative hypothesis that the HKGs tissue-weighted standard deviation tends to be lower than the standard deviation of all other proteins.

### Protein complex pairs (PCPs) curation process and benchmark

To evaluate the expected relationships between proteins within samples, we curated a list of protein-complex pairs (PCPs), defined as proteins mutually and exclusively co-expressed within protein complexes, therefore exhibiting similar expression values and/or high expression correlation. As a preliminary list, we curated PCPs from the CORUM database^27^. Then, within each dataset in our atlas, we calculated the Pearson correlation coefficient and mean absolute deviation (MAD) between each candidate PCP and filtered for PCPs with correlation coefficient > 0.9 and MAD<1. For each PCP, we counted in how many datasets it passed our correlation and MAD thresholds, and took the 50 PCPs with the highest count, provided in Supplementary.

We used two metrics to score the PCP benchmark; for the first metric, we calculated the mean absolute deviation between each PCP, by subtracting the expression of one protein from the other across all samples and taking the mean of the absolute values of the subtraction. We repeated this calculation for 10,000 randomly selected protein pairs and performed a Mann-Whitney U-test between the two distributions and calculated the p-value with the alternative hypothesis that the absolute deviation between PCPs tends to be lower than that of random pairs. For the second metric, we calculated the Pearson correlation coefficient between all PCPs, and determined the percentage of PCPs with a correlation coefficient > 0.7.

### Phenotypic markers curation process and differential expression (DE) benchmark

To evaluate the expected phenotypic variability within cancer indications, we compared two well defined breast cancer subtypes, Luminal A and Basal-like. To that end, through a literature survey, we manually cured a list of 64 proteins known to be overexpressed in either subtype, but not the other. The full list of phenotypic markers is provided in Supp. Table _. In this work, three breast cancer datasets were used for our harmonization effort (Krug et al.^28^, Asleh et al.^29^, Anurag et al.^30^). Therefore, we further curated the PAM50 annotations of these datasets’ tumor samples from their associated papers’ Supplementary clinical data tables.

To score this benchmark, and for each phenotypic marker, we performed a one-tailed Welch’s unequal variances t-test between Luminal A and Basal-like samples across the entire harmonized dataset, with the alternative hypothesis that the phenotypic marker’s mean is higher in its relevant subtype, and calculated its p-value. We then adjusted all p-values for multiple tests using the Benjamini-Hochberg False Discovery Rate method. Additionally, we calculated each marker’s fold change by subtracting its mean expression within its relevant PAM50 subtype from its mean expression within the other subtype. Finally, we calculated a “combined score” for each phenotypic marker with the following formula:

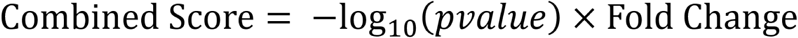

Last, we took the mean of the markers’ combined scores as the final score of this benchmark.

### Cross-batch technical replicates

To directly measure the success of protein-level harmonization, we took advantage of unique study designs in three datasets from our harmonized atlas. First, Hu et al.^14^ and McDermott et al.^15^ analyzed the same 99 ovarian cancer tumor samples, in two sets of technical replicates, at Johns Hopkins University (JHU) and the Pacific Northwest National Laboratory (PNNL), respectively. We considered these two sets as matching replicates. Second, Chowdhury et al.^16^, analyzed the same 49 ovarian cancer tumor samples in two separate batches of technical replicates; one batch consisted of Fresh-Frozen tumor samples, while the other batch consisted of formalin-fixed, paraffin-embedded (FFPE) samples of the same tumors. We further considered these two batches as another set of matching replicates.

To score the benchmark, we first filtered out noisy or flat-expression proteins. We calculated the inter-quantile range (IQR) of each protein within each dataset separately, by subtracting the expression at the 25th percentile from the expression at the 75th percentile, and considered only the top 50% proteins with the highest IQR within each batch for further analysis. Then, we calculated the protein-wise Pearson correlation coefficient between the two sets of replicates. Finally, we took the mean of the two resulting medians as this benchmark’s final score.

### Standardized Reanalysis of RAW Data

MzML files were analyzed using MSFragger v4.1^9,31,32,33,34,35^ against a Uniprot-reviewed protein FASTA with appended isoforms and decoy sequences. Strict trypsin digestion (cut after KR) was specified. Post-processing was performed with the Philosopher toolkit v5.1.1^36^, and statistical summarization and reporting were completed using TMT-Integrator v5.0.9. The search settings accounted for individual experimental conditions, including isotope error (−1/0/1/2/3), fixed modifications (Cysteine carbamidomethylation, Lysine TMT labeling), and variable modifications (Methionine oxidation, protein N-terminal acetylation, and TMT labeling of Serine and peptide N-termini). Searches were limited to tryptic peptides with up to two missed cleavages, a minimum of 15 peaks, precursor ion tolerance, and fragment tolerance set to 20 ppm, and a clear mz range of 125.5–131.5. Data was filtered using Philosopher’s sequential method, combining all pep.xml files. PSMs, peptides, and proteins were filtered to 1% FDR using the picked target-decoy strategy. MS1 intensity was extracted with IonQuant using a 10 ppm mass tolerance and 0.4 min retention time window. TMT-Integrator processed all PSM files together, with peptide-protein quantification set to both unique and razor peptides, and peptide-gene quantification set to unique peptides. Data normalization used median centering and an artificial reference channel.

Outlier samples, including those with poor or inconsistent peptide identification rates, low correlation with other samples, or significant deviations in intensity distributions, were excluded from further analysis. Additionally, dataset-level QC was performed, ensuring high protein coverage, minimal missing values, and consistency with the expected ratios between biological classes. Only data meeting these stringent QC criteria were retained for further analysis.

### Normalization

Sample-wise normalization was achieved by performing high-quantile (HQ) scaling, with the assumption that higher values in the sample-wise proteome-wide intensity distribution is typically measured with greater precision and, relatively, reflect a biologically invariant population of proteins, providing a reliable anchor for normalization. Therefore, following the standardized reanalysis of raw data, each sample was normalized by subtracting from all its protein expression intensity values the average of their 95th and 99th percentiles, with the following formula:

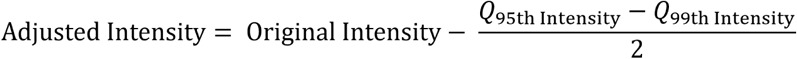

Additionally, we tested other normalization methods; Sample-wise and gene-wise Robust Scaling were achieved using the RobustScaler function from sklearn.preprocessing package with default parameters, on the entire original and transposed expression matrix, respectively. Sample-wise and gene-wise median scaling were performed by subtracting the sample-wise or gene-wise median from each value in the expression matrix, respectively.

### Tissue-wise batch effect removal and imputation

Missing values in proteomics datasets pose significant challenges for harmonization and batch effect correction methods. To address this, we developed a novel multi-step approach that leverages information from other datasets of the same tissue type for imputation, coupled with batch effect removal, thereby enhancing data completeness and consistency.

First, we applied HarmonizR^11^ an R package utilizing matrix dissection to implement ComBat batch effect correction without requiring prior imputation of missing values. Given that ComBat assumes datasets originate from the same underlying distribution, we applied it in a tissue-wise manner to account for biological variability inherent to different cancer types.

To accommodate intra-tissue heterogeneity, such as distinct molecular subtypes within a cancer type, we extended HarmonizR’s usage of ComBat within the “sva” R package^4^ to incorporate biological covariates (e.g., PAM50 subtypes in breast cancer). This modification allowed for more precise batch effect correction while preserving biologically meaningful differences.

Subsequently, we performed protein-wise left-tail imputation, estimating missing values for each protein (i) by sampling from a narrow normal distribution around the lower quantiles of the protein intensity distributions with the following formula:

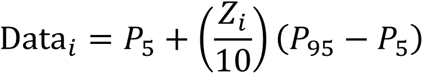

Where:

● Data_i_ is the value in the dataset at index i.
● P_5_ and P_95_ are the 5th and 95th percentiles of the data, respectively.
● Z_i_ is a standard normal random variable, Z_i_ ∼ N(0,1)

The term 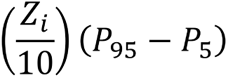 introduces a small random variation around the 5th percentile, scaled by the inter-percentile range between the 95th and 5th percentiles. This method imputes missing values by adding randomized noise to the lower end of the data distribution, ensuring that the imputed values remain within a plausible range. This step generated an initial complete reference matrix for each tissue type.

Dimensionality reduction was then conducted on the complete reference matrix using t-distributed stochastic neighbor embedding (t-SNE) to enhance local multidimensional similarities among samples. t-SNE was performed with perplexity 30, 100,000 iterations and on 3 principal components. In the reduced space, we calculated the pair-wise Euclidean distances between samples and identified the K-nearest neighbors (KNN) for each sample, where K=3. Finally, missing values in a sample were imputed by averaging the corresponding protein intensities from its KNN in other datasets, effectively replacing the left-tail estimates with more accurate imputations based on biological similarity. Any protein missing entirely from any individual tissue was subsequently dropped from further analysis.

### Autoencoder

To address residual batch effects and enhance integration across different tissue types, we employed an autoencoder neural network model. Autoencoders are well-suited for learning compressed representations of high-dimensional data, capturing essential features while minimizing noise.

In total, we trained well over a 1,000 models with a variety of different architectures and hyperparameter settings. The harmonization benchmarks guided us throughout this process, where eventually we manually selected the model with the best overall performance. The final autoencoder architecture consisted of an input layer matching the dimensionality of the proteomics data (8960 nodes), an encoder, a bottleneck layer, and a decoder. The encoder reduced the dimensionality of the input data using fully connected multilayer perceptrons with 2000, 1500, and 1000 nodes, while the decoder reconstructed the input data by reversing the compression process with layers of corresponding size. A bottleneck layer with 500 units served as the latent representation, capturing shared biological signals across datasets. To prevent overfitting, dropout regularization was applied at rates of 0.05 and 0.02 across the encoder and decoder layers, respectively. All layers used ReLU (Rectified Linear Unit) activation functions, except for the output layer, which employed a linear activation for reconstruction.

The training process was guided by two primary loss functions. The reconstruction loss, calculated as the mean absolute error (MAE) between the input and reconstructed data, ensured the preservation of essential biological features. Additionally, a consistency loss penalized deviations from the original protein-wise expression rankings of samples within each dataset, preserving dataset-wise relative rankings assumed to reflect true biological variability. The consistency loss was calculated as the mean squared error (MSE) of the pair-wise distances between the input and reconstructed ranks, for each protein and within each dataset separately. The combined loss function was defined as:

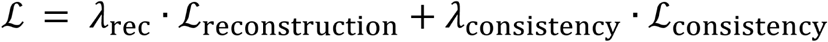

Where

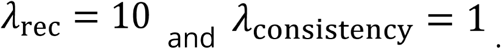

Optimization was performed using the Adam optimizer, which is well-suited for non-convex optimization in neural networks, with moving mean and variance exponential decays of 0.5 and 0.9, respectively. The learning rate for the reconstruction task was set to 0.0001. Exponential learning rate decay with a decay factor of 0.9 was applied every 64 steps to stabilize convergence. A weight decay of 1e−05 was used as additional regularization to prevent overfitting. Gradients were clipped to a maximum norm of 1.0 to ensure stable training dynamics.

The model was implemented using the PyTorch framework (v2.2.2-cuda11.8)^37^. A batch size of 500 was set for 650 epochs, without early stopping.

### Algorithm evaluation

Throughout the development of our algorithm, we employed our novel harmonization benchmarks (described above) to assess the success of different harmonization steps. Before the imputation step, missing values were included in our benchmark analysis. The comparison to the pan-cancer works of Li et al.^17^ and Zhang et al.^18^ were also performed with missing values. Conversely, to enable a fair and robust comparison between other harmonization methods and our algorithm, we implemented a base-line imputation process before applying the different methods. As previously described, we performed a protein-wise left-tail imputation, estimating missing values for each protein by sampling from a narrow normal distribution around the lower quantiles of the protein intensity distributions.

For comparison, the removeBatchEffect function from the “Limma” package^5^ (https://rdrr.io/bioc/limma/man/removeBatchEffect.html) was run in R, with default parameters. The ComBat function from the “sva” package^4^ was also run in R with default parameters.

### Statistical data analysis and visualization

#### Pan-Cancer Indication Expansion and Prioritization

A list of oncologic therapeutic targets and their clinically approved indications, aimed to reflect the most prominent and well-established therapies, were manually curated from literature. We then calculated the tissue-weighted median of each target, and z-score transformed the values per target, and target-tissue pairs with expression above the tissue-weighted median were annotated as clinically significant. Precision and recall were calculated by calling “precision_score” and “recall_score” functions from the sklearn.metrics package against all significant and non-significant target-tissue pairs with default parameters. Visualization was performed with the “scatterplot” function from the Seaborn package and a “RdBu_r” color palette.

#### Target Discovery for Pan-Cancer Subtypes

Pathway enrichment scores were calculated by calling the “gsva” function from the gseapy package on the integrated pan-cancer expression matrix with the “KEGG_2021_Human” library, min_size=15 and max_size=500. Resulting pathways were then manually filtered for oncology relevance. Hierarchical clustering and clustermap visualization was performed by calling the “clustermap” function from the Seaborn package on the pathway enrichment scores matrix with a “coolwarm” color map. PCA was calculated with the “PCA” function from the sklearn.decomposition package with default parameters, and visualized with Seaborn “scatterplot”. Samples were assigned to clusters C1-C3 by their respective tissues according to the visualization. Differentially expressed proteins for each cluster were selected by calculating the fold change for each protein between each cluster and the other clusters. Sample mutation data was curated as previously described. Clinically relevant targets were determined by literature review. Druggability data was acquired from OpenTargets^38^ website.

#### Consistent Prognostic Biomarker Discovery for Lung Adenocarcinoma

We combined two independent LUAD datasets (Lehtiö et al.^21^ and Gilette et al.^22^) as our discovery cohort. Sample patients’ overall survival was collected as previously described, and split into good (OS > 3) and poor (OS < 2) prognosis groups. Multiple ANOVA was performed by calling the “SelectKBest” function from the sklearn.feature_selection package with score_func=f_classif and k=“all”. In the iterative feature selection step, 80% and 20% were randomly assigned for the train and test sets, respectively, for every feature combination. For the logistic regression classifier, we used “LogisticRegression” from the sklearn.linear_model package with max_iter=1000. The ROC AUC was calculated by calling the “roc_auc_score” function from the sklearn.metrics package. The sample-wise prognostic biomarker heatmap was visualized with the “pheatmap” function in R. The functional protein association network was generated and visualized with STRING^39^, providing the top 20 differentially expressed proteins between the two prognostic groups as input.

## Acknowledgments

We acknowledge Gali Arad and Anjana Shenoy from Protai Bio for their valuable contributions to the writing process, insightful feedback, and thorough review of the manuscript.

## References

1. Phua, Ser-Xian, Kai-Peng Lim, and Wilson Wen-Bin Goh. “Perspectives for Better Batch Effect Correction in Mass-Spectrometry-Based Proteomics.” Computational and Structural Biotechnology Journal, vol. 20, 2022, pp. 4369–4375. 10.1016/j.csbj.2022.08.022

2. Zhang, Xu et al. “Assessing the impact of protein extraction methods for human gut metaproteomics.” Journal of proteomics vol. 180 (2018): 120–127. doi:10.1016/j.jprot.2017.07.001

3. Danko, Katerina, et al. “Comparative Analysis of Methods for Batch Correction in Proteomics — a Two-Batch Case”. Biological Communications, vol. 68, no. 1, May 2023, pp. 56–61, doi:10.21638/spbu03.2023.106.

4. Leek, J. T., et al. sva: Surrogate Variable Analysis. R package version 3.50.0, 2023, Bioconductor, doi:10.18129/B9.bioc.sva.

5. Ritchie, M.E., Phipson, B., Wu, D., Hu, Y., Law, C.W., Shi, W., and Smyth, G.K. (2015). limma powers differential expression analyses for RNA-sequencing and microarray studies. Nucleic Acids Research 43(7), e47.

6. Edwards, Nathan J., et al. “The CPTAC Data Portal: A Resource for Cancer Proteomics Research.” Journal of Proteome Research, vol. 14, no. 6, American Chemical Society, 5 June 2015, pp. 2707–2713. 10.1021/pr501254j.

7. Rong, Zhiwei, et al. “NormAE: Deep Adversarial Learning Model to Remove Batch Effects in Liquid Chromatography Mass Spectrometry-Based Metabolomics Data.” Analytical Chemistry, vol. 92, no. 7, 2020, pp. 5082–5090. American Chemical Society, 10.1021/acs.analchem.9b05460.

8. Dai, Chengxin, et al. “quantms: a cloud-based pipeline for quantitative proteomics enables the reanalysis of public proteomics data.” Nature Methods 21.9 (2024): 1603–1607.

9. Kong, A. T., Leprevost, F. V., Avtonomov, D. M., Mellacheruvu, D., & Nesvizhskii, A. I. (2017). MSFragger: ultrafast and comprehensive peptide identification in mass spectrometry–based proteomics. Nature Methods, 14(5), 513–520.

10. Välikangas T, Suomi T, Elo LL. A systematic evaluation of normalization methods in quantitative label-free proteomics. Brief Bioinform. 2018;19(1):1–11.

11. Voß, H., et al. “HarmonizR enables data harmonization across independent proteomic datasets with appropriate handling of missing values. Nat Commun. 2022; 13 (1): 3523.”

12. Zhou, Y., Lih, T.M., Pan, J. et al. Proteomic signatures of 16 major types of human cancer reveal universal and cancer-type-specific proteins for the identification of potential therapeutic targets. J Hematol Oncol 13, 170 (2020). 10.1186/s13045-020-01013-x

13. Song, Q., Yang, Y., Jiang, D. et al. Proteomic analysis reveals key differences between squamous cell carcinomas and adenocarcinomas across multiple tissues. Nat Commun 13, 4167 (2022). 10.1038/s41467-022-31719-0

14. Hu, Yingwei et al. “Integrated Proteomic and Glycoproteomic Characterization of Human High-Grade Serous Ovarian Carcinoma.” Cell reports vol. 33,3 (2020): 108276. doi:10.1016/j.celrep.2020.108276

15. McDermott, Jason E et al. “Proteogenomic Characterization of Ovarian HGSC Implicates Mitotic Kinases, Replication Stress in Observed Chromosomal Instability.” Cell reports. Medicine vol. 1,1 (2020): 100004. doi:10.1016/j.xcrm.2020.100004

16. Chowdhury, Shrabanti et al. “Proteogenomic analysis of chemo-refractory high-grade serous ovarian cancer.” Cell vol. 186,16 (2023): 3476–3498.e35. doi:10.1016/j.cell.2023.07.004

17. Li, Yize, et al. “Proteogenomic data and resources for pan-cancer analysis.” Cancer cell 41.8 (2023): 1397–1406.

18. Zhang, Yiqun, et al. “Proteogenomic characterization of 2002 human cancers reveals pan-cancer molecular subtypes and associated pathways.” Nature Communications 13.1 (2022): 2669.

19. Yan, Ge, et al. “Relationship between EGFR expression and subcellular localization with cancer development and clinical outcome.” Oncotarget [Online], 10.20 (2019): 1918–1931. Web. 9 Feb. 2025

20. Hu, Beiyuan et al. “Inhibition of EGFR Overcomes Acquired Lenvatinib Resistance Driven by STAT3-ABCB1 Signaling in Hepatocellular Carcinoma.” Cancer research vol. 82,20 (2022): 3845–3857. doi:10.1158/0008-5472.CAN-21-4140

21. Lehtiö, Janne, et al. “Proteogenomics of Non-Small Cell Lung Cancer Reveals Molecular Subtypes Associated with Specific Therapeutic Targets and Immune-Evasion Mechanisms.” Nature Cancer, vol. 2, no. 11, 2021, pp. 1224–1242, doi:10.1038/s43018-021-00259-9.

22. Gillette, Michael A et al. “Proteogenomic Characterization Reveals Therapeutic Vulnerabilities in Lung Adenocarcinoma.” Cell vol. 182,1 (2020): 200–225.e35. doi:10.1016/j.cell.2020.06.013

23. Soltis, Anthony R., et al. “Proteogenomic Analysis of Lung Adenocarcinoma Reveals Tumor Heterogeneity, Survival Determinants, and Therapeutically Relevant Pathways.” Cell Reports Medicine, vol. 3, no. 11, 15 Nov. 2022, Elsevier, 10.1016/j.xcrm.2022.100819.

24. Ratna R. Thangudu, Michael Holck, Deepak Singhal, Alexander Pilozzi, Nathan Edwards, Paul A. Rudnick, Marcin J. Domagalski, Padmini Chilappagari, Lei Ma, Yi Xin, Toan Le, Kristen Nyce, Rekha Chaudhary, Karen A. Ketchum, Aaron Maurais, Brian Connolly, Michael Riffle, Matthew C. Chambers, Brendan MacLean, Michael J. MacCoss, Peter B. McGarvey, Anand Basu, John Otridge, Esmeralda Casas-Silva, Sudha Venkatachari, Henry Rodriguez, Xu Zhang; NCI’s Proteomic Data Commons: A Cloud-Based Proteomics Repository Empowering Comprehensive Cancer Analysis through Cross-Referencing with Genomic and Imaging Data. Cancer Research Communications 1 September 2024; 4 (9): 2480–2488.. 10.1158/2767-9764.CRC-24-0243 https://proteomic.datacommons.cancer.gov/pdc/

25. Deutsch EW, Bandeira N, Perez-Riverol Y, Sharma V, Carver JJ, Mendoza L, Kundu DJ, Wang S, Bandla C, Kamatchinathan S, Hewapathirana S, Pullman BS, Wertz J, Sun Z, Kawano S, Okuda S, Watanabe Y, MacLean B, MacCoss MJ, Zhu Y, Ishihama Y, Vizcaíno JA. The ProteomeXchange consortium at 10 years: 2023 update. Nucleic Acids Res. 2023 Jan 6;51(D1):D1539–D1548. doi: 10.1093/nar/gkac1040. PMID: 36370099; PMCID: PMC9825490.

26. Perez-Riverol Y, Bandla C, Kundu DJ, Kamatchinathan S, Bai J, Hewapathirana S, John NS, Prakash A, Walzer M, Wang S, Vizcaíno JA. The PRIDE database at 20 years: 2025 update. Nucleic Acids Res. 2024 Nov:gkae1011. doi: 10.1093/nar/gkae1011. PMID: 39494541.

27. Tsitsiridis, George et al. “CORUM: the comprehensive resource of mammalian protein complexes-2022.” Nucleic acids research vol. 51,D1 (2023): D539–D545. doi:10.1093/nar/gkac1015

28. Krug, Karsten, et al. “Proteogenomic Landscape of Breast Cancer Tumorigenesis and Targeted Therapy.” Cell, vol. 183, no. 5, 25 Nov. 2020, pp. 1436–1456.e31, Elsevier, doi:10.1016/j.cell.2020.10.036

29. Asleh, K., Negri, G.L., Spencer Miko, S.E. et al. Proteomic analysis of archival breast cancer clinical specimens identifies biological subtypes with distinct survival outcomes. Nat Commun 13, 896 (2022). 10.1038/s41467-022-28524-0

30. Anurag, Meenakshi, et al. “Proteogenomic Markers of Chemotherapy Resistance and Response in Triple-Negative Breast Cancer.” Cancer Discovery, vol. 12, no. 11, American Association for Cancer Research, 2 Nov. 2022, pp. 2586–2605, 10.1158/2159-8290.CD-22-0200

31. Yu, F., Teo, G. C., Kong, A. T., Haynes, S. E., Avtonomov, D. M., Geiszler, D. J., & Nesvizhskii, A. I. (2020). Identification of modified peptides using localization-aware open search. Nature Communications, 11(1), 1–9.

32. Polasky, D. A., Yu, F., Teo, G. C., & Nesvizhskii, A. I. (2020). Fast and Comprehensive N-and O-glycoproteomics analysis with MSFragger-Glyco. Nature Methods, 17(11), 1125–1132.

33. Yu, F., Haynes, S. E., Teo, G. C., Avtonomov, D. M., Polasky, D. A., & Nesvizhskii, A. I. (2020). Fast Quantitative Analysis of timsTOF PASEF Data with MSFragger and IonQuant. Molecular & Cellular Proteomics, 19(9), 1575–1585.

34. Polasky, D., Geiszler, D., Yu, F., Li, K., Teo, G. C., & Nesvizhskii, A. I., (2023). MSFragger-Labile: A flexible method to improve labile PTM analysis in proteomics. Molecular & Cellular Proteomics, 22(5), 100538.

35. Yu, F., Teo, G. C., Kong, A. T., Fröhlich, K., Li, G. X., Demichev, V., & Nesvizhskii, A. I., (2023). Analysis of DIA proteomics data using MSFragger-DIA and FragPipe computational platform. Nature Communications, 14, 4154.

36. da Veiga Leprevost F, Haynes SE, Avtonomov DM, Chang HY, Shanmugam AK, Mellacheruvu D, Kong AT, Nesvizhskii AI. Philosopher: a versatile toolkit for shotgun proteomics data analysis. Nat Methods. 2020 Sep;17(9):869–870. doi: 10.1038/s41592-020-0912-y. PMID: 32669682; PMCID: PMC7509848.

37. Ansel, J., Yang, E., He, H., Gimelshein, N., Jain, A., Voznesensky, M., Bao, B., Bell, P., Berard, D., Burovski, E., Chauhan, G., Chourdia, A., Constable, W., Desmaison, A., DeVito, Z., Ellison, E., Feng, W., Gong, J., Gschwind, M., Hirsh, B., Huang, S., Kalambarkar, K., Kirsch, L., Lazos, M., Lezcano, M., Liang, Y., Liang, J., Lu, Y., Luk, C., Maher, B., Pan, Y., Puhrsch, C., Reso, M., Saroufim, M., Siraichi, M. Y., Suk, H., Suo, M., Tillet, P., Wang, E., Wang, X., Wen, W., Zhang, S., Zhao, X., Zhou, K., Zou, R., Mathews, A., Chanan, G., Wu, P., & Chintala, S. (2024). PyTorch 2: Faster Machine Learning Through Dynamic Python Bytecode Transformation and Graph Compilation [Conference paper]. 29th ACM International Conference on Architectural Support for Programming Languages and Operating Systems, Volume 2 (ASPLOS ’24). 10.1145/3620665.3640366

38. Buniello, Annalisa, et al. “Open Targets Platform: Facilitating Therapeutic Hypotheses Building in Drug Discovery.” Nucleic Acids Research, vol. 53, no. D1, 6 Jan. 2025, pp. D1467–D1475, 10.1093/nar/gkae1128.

39. Szklarczyk, Damian, et al. “The STRING Database in 2021: Customizable Protein–Protein Networks, and Functional Characterization of User-Uploaded Gene/Measurement Sets.” Nucleic Acids Research, vol. 49, no. D1, 2021, pp. D605–D612, doi:10.1093/nar/gkaa1074.

